# Inclusion of Variants Discovered from Diverse Populations Improves Polygenic Risk Score Transferability

**DOI:** 10.1101/2020.05.21.108845

**Authors:** Taylor B. Cavazos, John S. Witte

## Abstract

The majority of polygenic risk scores (PRS) have been developed and optimized in individuals of European ancestry and may have limited generalizability across other ancestral populations. Understanding aspects of PRS that contribute to this issue and determining solutions is complicated by disease-specific genetic architecture and limited knowledge of sharing of causal variants and effect sizes across populations. Motivated by these challenges, we undertook a simulation study to assess the relationship between ancestry and the potential bias in PRS developed in European ancestry populations. Our simulations show that the magnitude of this bias increases with increasing divergence from European ancestry, and this is attributed to population differences in linkage disequilibrium and allele frequencies of European discovered variants, likely as a result of genetic drift. Importantly, we find that including into the PRS variants discovered in African ancestry individuals has the potential to achieve unbiased estimates of genetic risk across global populations and admixed individuals. We confirm our simulation findings in an analysis of HbA1c, asthma, and prostate cancer in the UK Biobank. Given the demonstrated improvement in PRS prediction accuracy, recruiting larger diverse cohorts will be crucial—and potentially even necessary—for enabling accurate and equitable genetic risk prediction across populations.

## INTRODUCTION

Increasing research into polygenic risk scores (PRS) for disease prediction highlights their clinical potential for informing screening, therapeutics, and lifestyle^1^. While their use enables risk prediction in individuals of European ancestry, PRS can have widely varying and much lower accuracy when applied to non-European populations^2–4^. Although the nature of this bias is not well understood, it can be attributed to the vast overrepresentation of European ancestry individuals in genome-wide association studies (GWAS), which is 4.5-fold higher than their percentage of the world population; conversely, there is underrepresentation of diverse populations such as individuals of African ancestry in GWAS, which is one fifth their percentage^3^. Potential explanations for the limited portability of European derived PRS across populations includes differences in population allele frequencies and linkage disequilibrium, the presence of population-specific causal variants or effects, or potential differences in gene-gene or gene-environment interactions^4^. However, in traits such as body mass index and type 2 diabetes, 70 to 80% of European-based PRS accuracy loss in African ancestry has been attributed to differences in allele frequency and linkage disequilibrium; therefore, most causal variants discovered in Europeans are likely to be shared^5^. Recent methods developed to improve PRS accuracy in non-Europeans have prioritized the use of European discovered variants and population specific weighting^6–8^. However, only small gains in accuracy are possible with limited sample sizes of non-European cohorts^4^.

PRS have been applied and characterized within global populations, but there is limited understanding of PRS accuracy in recently admixed individuals and whether this varies with ancestry. Studies applying PRS in diverse populations^3–5,9^ or exploring potential statistical approaches to improve accuracy in such populations^6,10^ typically present performance metrics averaged across all admixed individuals. Only one study to date has suggested that PRS accuracy may be a function of genetic admixture (i.e., for height in the UK Biobank^8^). However, it is unknown if the relationship between accuracy and ancestry exists when variants are discovered in non-European populations or what the best approach for applying PRS to admixed individuals will be when there are adequately powered GWAS in non-European populations.

To help answer these questions, here we systematically and empirically explore the relationship between PRS performance and ancestry within African, European, and admixed ancestry populations through simulations. We highlight PRS building approaches that will achieve unbiased estimates across global populations and admixed individuals with future recruitment and representation of non-European ancestry individuals in GWAS. We also investigate reasons for loss of PRS accuracy, and attribute this to population differences in linkage disequilibrium (LD) tagging of causal variants by lead GWAS variants, as well as allele frequency biases potentially due to genetic drift undergone by European ancestry populations. Finally, we confirm our simulation findings by application to data on HbA1c levels, asthma, and prostate cancer in individuals of European and individuals of African ancestry from the UK Biobank.

## MATERIAL AND METHODS

### Simulation of Population Genotypes

We used the coalescent model (msprime v.7.3^11^) to simulate European (CEU) and African (YRI) genotypes, based on whole-genome sequencing of HapMap populations, for chromosome 20 as described previously by Martin et al.^2^ Genotypes were modeled after the demographic history of human expansion out of Africa^12^, assuming a mutation rate of 2 x 10^−8^. We simulated 200,000 Europeans and 200,000 Africans for each simulation trial, for a total of 50 independent simulations (20 million total individuals). We generated founders from an additional 1,000 Europeans and 1,000 Africans (10,000 total across the 50 simulations) to simulate 5,000 admixed individuals (250,000 total across the 50 simulations) with RFMIX v.2^13^ assuming two-way admixture between Europeans and Africans with random mating and 8 generations of admixture.

### True and GWAS Estimated Polygenic Risk Scores

We generated true genetic liability for all European, African, and admixed individuals within each simulation trial^2^. Briefly, m variants evenly spaced throughout the simulated genotypes were selected to be causal and the effect sizes (*β*) were drawn from a normal distribution 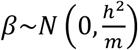, where h^2^ is the heritability^2^. Constant heritability and complete sharing of effect sizes in African ancestry and European ancestry individuals was assumed. The true genetic liability was computed as the summation of all variant effects multiplied by their genotype for each individual 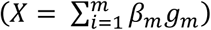 standardized to ensure total variance of 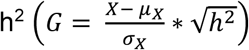. Finally, the non-genetic effect (*ε* = *N*(0, 1 − *h*^2^)) standardized to explain the remainder of the phenotypic variation 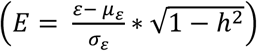 was added to the genetic risk defining the total trait liability (*G* + *E*)^2^. Cases were selected from the extreme tail of the liability distribution, assuming a 5% disease prevalence. An equal number of controls and 5,000 testing samples were randomly selected from the remainder of the distribution; all 5,000 admixed individuals were also used for testing. Across simulation replicates we varied causal variants (m = {200, 500, 1000}) and trait heritability (h ^2^ = {0.33, 0.50, 0.67}); however, for simplicity main text results assume m = 1000 and h^2^ = 0.50.

The estimated PRS were constructed from GWAS of the simulated genotypes (modeled after chromosome 20) in European and African ancestry populations, each with 10,000 cases and 10,000 controls. Odds ratios (ORs) were estimated for all variants with minor allele frequency (MAF) > 1% and statistical significance of association was assessed with a chi-squared test. While causal variants may be included in the estimated PRS, they are drawn from the total allele frequency spectrum; thus, they are primarily rare (93% and 87% of causal variants have MAF < 1% in European and African ancestry populations when m = 1000) and restricted from our analysis. For each population, variants were selected for inclusion into the estimated PRS by p-value thresholding (p = 0.01 (*Main Text*), 1×10^−4^, and 1×10^−6^ (*Supplements*)) and clumping (r^2^ < 0.2) in a 1 Mb window within the GWAS population, where r^2^ is the squared Pearson correlation between pairs of variants. A fixed-effects meta-analysis was also performed to calculate the inverse-variance weighted average of the ORs in African and European ancestry populations, and LD r^2^ values for clumping used both datasets as the reference.

For each individual, an estimated PRS was calculated as the sum of the log(OR) (i.e., the PRS ‘weights’) multiplied by their genotype for all independent and significant variants at a given threshold. The PRS were constructed for testing samples with variants and weights each selected from European or African ancestry GWAS, or a fixed-effects meta of both combined. Additional multi-ancestry PRS approaches^7,10^ were also explored for admixed individuals. Accuracy was measured by Pearson’s correlation (*r*) between the true genetic liability and estimated PRS within each population. Across simulation trials, averages and ninety-five percent confidence intervals for *r* were calculated following a Fisher z-transformation for approximate normality^14^. The statistical significance of differences in accuracy between PRS approaches was assessed within ancestry groups, defined by global genome-wide European ancestry proportions, with a z-test (also following Fisher transformation). Specifically, within each simulation trial the z-statistic, measuring the difference between two PRS approaches, was computed and a two-sided p-value was obtained; results were summarized across trials by taking the median p-value. While using *r* as a measure of accuracy has the added benefit of being independent from heritability, in admixed individuals we also calculate the proportion of variance (R^2^) for total trait liability (genetic and environmental component) explained by the estimated PRS.

### Multi-ancestry PRS

#### Local Ancestry Weighting PRS

In addition to genotypes of simulated admixed individuals, RFMIX^13^ also outputs the local ancestry at each locus for every individual. Thus, we used this information to create a local ancestry weighted PRS that is not impacted by ancestry inference errors. For admixed African and European ancestry individuals an ancestry-specific PRS was constructed for each population (k) by limiting each PRS to variants found in that ancestry-specific subset of the genome (*i* ∈ *k*), as defined by local ancestry, and weighting using variant effects discovered in that population^7^. Each ancestry-specific PRS was then combined, weighted by the genome-wide global ancestry proportion (*ρ*_*k*_) for that individual as follows^7^:

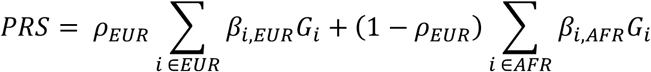

In this way each individual has a PRS constructed from the same independent variants with personalized weights that are unique to the individual’s local ancestry.

#### Linear Mixture of Multiple Ancestry-Specific PRS

Using a linear mixture approach developed by Márquez-Luna et al.^10^ we combined two PRS estimated in each of our global training populations

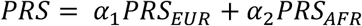

where individual PRS were constructed using significant and independent variants (p < 0.01 and r^2^ < 0.2 in a 1Mb window) and effect sizes from a GWAS in that training population. For simulations, mixing weights (*α*_1_ and *α*_2_) were estimated in an independent African ancestry testing population and as validation, accuracy was assessed in our simulated admixed ancestry individuals.

### Application to Real Data

We obtained genome-wide summary statistics for HbA1c^15^, asthma^16,17^, and prostate cancer^18,19^ calculated in European and African ancestry individuals (Table S1). Summary statistic variants that were not present in both the UK Biobank European and African ancestry testing populations were removed. PRS for each phenotype were constructed from associated and independent GWAS variants within each training population by p-value thresholding (p= {5×10^−8^, 1×10^−7^, 5×10^− 7^, 1×10^−6^, 5×10^−6^, 1×10^−5^, 5×10^−5^, 1×10^−4^, 5×10^−4^, 1×10^−3^, 5×10^−3^, 0.01, 0.05, 0.1, 0.5, 1}) and clumping (LD r^2^ < 0.2) of variants within 1Mb with PLINK^20^. Additionally a fixed-effects meta-analysis of the two populations was performed using METASOFT v2.0.1^21^. The selected PRS variants exhibited limited heterogeneity between the European and African ancestry training set summary statistics. In particular, of all possible European (African) ancestry selected PRS variants, only 5.4% (9.4%), 6.9% (5.7%), and 7.0% (4.8%) were heterogeneous between the two groups for HbA1c, asthma, and prostate cancer, respectively (i.e., I^2^ > 25% and Q p-value < 0.05).

PRS performance was evaluated in an independent cohort using genotype and phenotype data for individuals of European ancestry and individuals of African ancestry (Table S1) from the UK Biobank, imputation and quality control previously described^22^. We undertook extensive post-imputation quality control of the UK Biobank, including the exclusion of relatives and ancestral outliers from within each group. Specifically, analyses were limited to self-reported European and African ancestry individuals, with additional samples excluded if genetic ancestry PCs did not fall within five standard deviations of the self-reported population mean. For each individual, their PRS was computed as the weighted sum of the genotype estimates of effect on each phenotype from the discovery studies (Table S1), multiplied by the genotype at each variant. For each population-specific variant set, weights from either the European or African summary statistics or the fixed-effects meta-analysis were used. A total of 96 polygenic risk scores were evaluated in each phenotype exploring the impact of ancestral population (two scenarios), p-value threshold (16 scenarios), and variant weighting (three scenarios). The proportion of variation explained by each PRS (partial-R^2^) approach was assessed for UKB European-ancestry and African-ancestry individuals separately. The partial-R^2^ was calculated from the difference in R^2^ values following linear regression of HbA1c levels on age, sex, BMI, and PCs (1-10) with and without the PRS also included. Similarly, for asthma and prostate cancer, we determined the Nagelkerke’s pseudo partial-R^2^ following logistic regression of case status on age, sex (asthma only), BMI (prostate cancer only), and PCs (1-10) with and without the PRS. Additionally, in African ancestry individuals we created a combined PRS (*α*_1_*PRS*_*EUR*_ + *α*_2_*PRS*_*AFR*_) where *PRS*_*EUR*_ and *PRS*_*AFR*_ was the most optimal PRS using variants from the designated population and the weight and p-value that resulted in the highest accuracy; albeit in sample, optimization was done within a single PRS to ensure limited overfitting of the combined model^10^. We used 5-fold cross validation to assess model performance in which 80% of the cohort was used to estimate the mixing coefficients (*α*_1_ and *α*_2_) and the combined PRS partial-R^2^ was calculated in the remaining 20% of the data. Reported partial-R^2^ was averaged across folds^10^. For our binary phenotypes with unbalanced cases and controls we used stratified 5-fold cross validation.

## RESULTS

### Generalizability of European Derived Risk Scores to an Admixed Population

We constructed PRS from our simulated European datasets and applied them to independent simulated European, African, and admixed testing populations, assuming 1000 true causal variants (m) and trait heritability (h^2^) of 0.5. On average, 1552 (range = [1134-1920]) variants were selected for inclusion into the PRS at p-value < 0.01 and LD r^2^ < 0.2 (Table 1). The average accuracy across replicates (50 simulations), measured by the correlation (*r*) between the true and inferred genetic risk, was much higher when applying the PRS to Europeans (*r* = 0.77; 95% CI = [0.76, 0.77]) than to Africans (*r* = 0.45; 95% CI = [0.44, 0.47]; Figure 1). This is in agreement with decreased performance seen in real data when applying a European derived genetic risk score to an African population^2–5^.

**Table 1.** Summary of PRS Variants and Causal Tagging across Simulations. The set of PRS variants from each GWAS and the fixed-effects meta-analysis were selected by p-value thresholding (p < 0.01) and clumping (r^2^ < 0.2) across the 50 simulations. Each PRS variant was compared to the causal set of variants (m = 1000) within each simulation to determine the direct overlap between the two sets and the LD r^2^ between the PRS variant and every causal variant within a 1000 kb window. The total number of selected PRS variants that tag at least one cau sal variant at r^2^ greater than 0.8, 0.6, 0.4, or 0.2 is listed in the table.

| GWAS Population | Total # PRS<br>Variants (p<0.01) | # Causal | # in LD with a Causal Variant |  |  |  |
| --- | --- | --- | --- | --- | --- | --- |
| | | | $r^2 > 0.8$ | $r^2 > 0.6$ | $r^2 > 0.4$ | $r^2 > 0.2$ |
| European | 1552 [1134-1920] | 18 [10-26] |  |  |  |  |
| LD in Europeans |  |  | 27 [16-40] | 32 [22-44] | 39 [25-55] | 58 [38-80] |
| LD in Africans |  |  | 20 [9-36] | 25 [16-42] | 34 [24-54] | 53 [35-70] |
| African | 5269 [4462-6071] | 28 [18-40] | – | – | – | – |
| LD in Europeans |  |  | 94 [67-122] | 132 [95-171] | 183 [123-238] | 280 [202-364] |
| LD in Africans |  |  | 37 [26-48] | 48 [34-61] | 67 [50-89] | 127 [81-170] |
| Fixed-Effects Meta | 92 [38-197] | 12 [5-22] | – | – | – | – |
| LD in Europeans |  |  | 15 [6-26] | 17 [6-28] | 21 [9-39] | 29 [16-47] |
| LD in Africans |  |  | 13 [6-21] | 14 [6-25] | 17 [9-29] | 24 [10-43] |

\* The number of variants is reported as the average and range [low-high] across the 50 simulations

**Figure 1.**
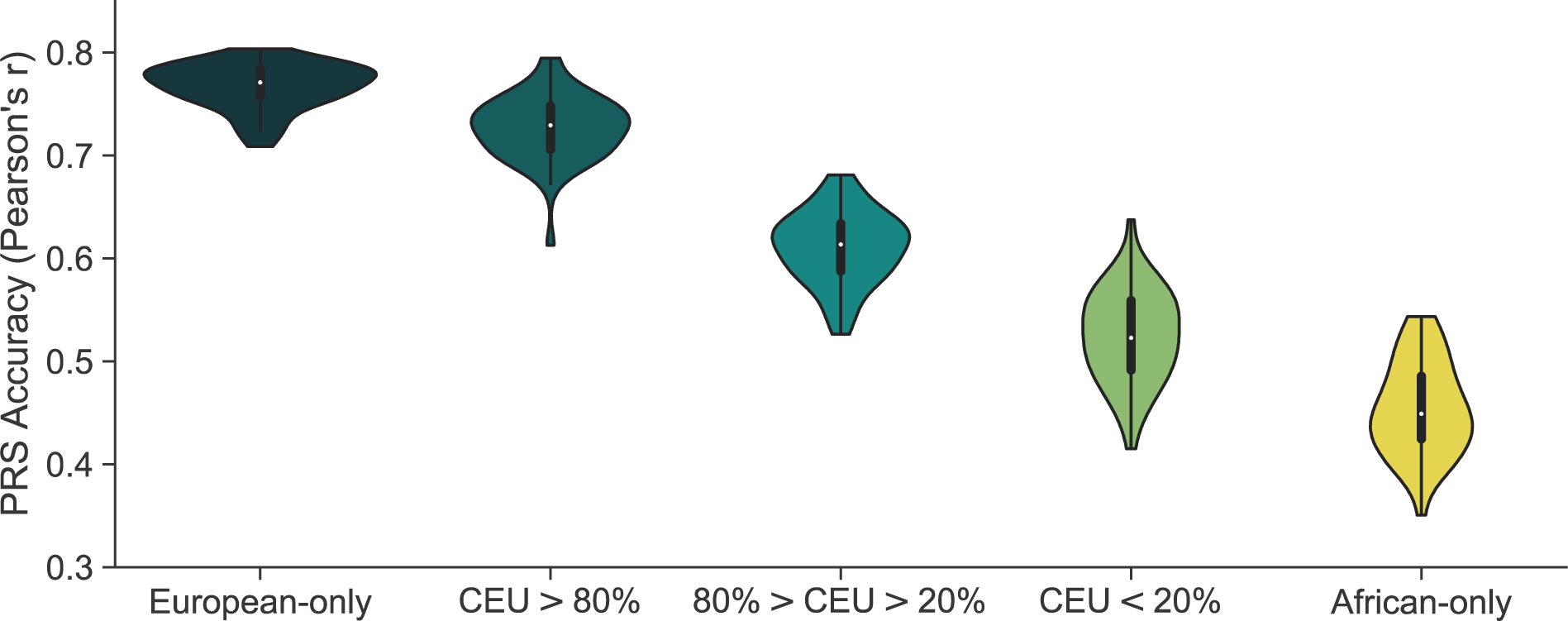
Accuracy of European Derived PRSs by Proportion of Total Ancestry. **Accuracy of PRS, with variants and weights from a European GWAS, decreases linearly with increasing proportion of African ancestry**. Variants and weights were extracted from a GWAS of 10000 European cases and 10000 European controls. PRS accuracy was computed as the Pearson’s correlation between the true genetic risk and GWAS estimated risk score across 50 simulations in independent test populations of 5000 Europeans, 5000 Africans, and 5000 admixed individuals. Admixed individuals were grouped based on their proportion of genome-wide European ancestry. Simulations assume 1000 causal variants and a heritability of 0.5 to compute the true genetic risk. A p-value of 0.01 and LD r^2^ cutoff of 0.2 was used to select variants for the estimated risk score.

To understand the relationship between ancestry and PRS accuracy, admixed individuals were stratified by their proportion of genome-wide European (CEU) ancestry: high (100%>CEU>80%), intermediate (80%>CEU>20%), and low (20%>CEU>0%). PRS performance decreased with decreasing European ancestry (Figure 1). Average accuracy (Pearson’s correlation) for the high, intermediate, and low European ancestry groups was 0.73 (95% CI = [0.72, 0.74]), 0.61 (95% CI = [0.60, 0.62]), and 0.53 (95% CI = [0.51, 0.54]), respectively (Figure 1). In comparison to Europeans, the performance of the European derived PRS was significantly lower in individuals with intermediate (20% decrease, *p* = 1.27×10^−47^), and low (32% decrease, *p* = 6.48×10^−16^) European ancestry, and with African-only ancestry (41% decrease, *p* = 8.00×10^−155^). There was no significant difference for individuals with high (5.3% decrease, *p* = 0.09) European ancestry. These trends remained consistent when varying the genetic architecture (Figure S1), specifically decreasing the number of causal variants (m) and varying the trait heritability (h^2^). Additionally, the relationship between ancestry and accuracy persisted with the inclusion of variants at lower p-value thresholds (Figure S2).

By further binning admixed individuals into deciles of global European ancestry and determining the variance explained of the total liability (genetics and environment) by the PRS, we estimated a 1.34% increase in accuracy for each 10% increase in global European ancestry, replicating a previous analysis of height in the UK Biobank^8^. The slope of this linear relationship increased with increasing heritability but was not found to vary with the number of true causal variants (Figure S3).

### Population Specific Weighting of European Selected Variants

Using a well-powered GWAS from our simulated African cohort (10,000 cases and 10,000 controls), we aimed to explore the potential accuracy gains achieved from a PRS with European selected variants, but with population specific weighting of these variants. We applied three different weighting approaches to incorporate non-European effect sizes: (1) effect sizes from an African ancestry GWAS for all variants; (2) effect sizes from a fixed-effects meta-analysis of European and African ancestry GWAS for all variants, both having 10,000 cases and 10,000 controls; and (3) population specific weights based on the local ancestry for an individual at each variant in the PRS (Figure 2).

**Figure 2.**
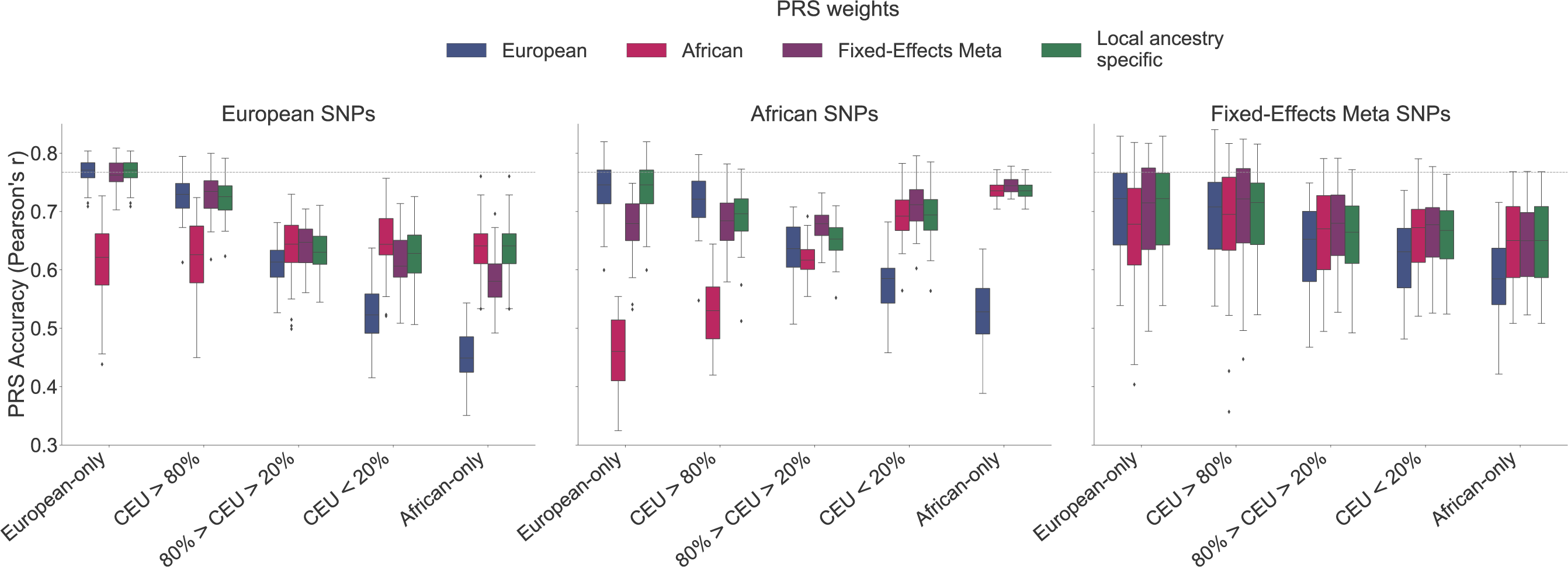
PRS Construction Approaches and Performance in Admixed Individuals. **Using significant variants from an African Ancestry GWAS with population-specific weights results in less disparity in PRS accuracy across populations**. PRS were constructed using variants and weights selected from either a European or African population (10000 cases, 10000 controls each) or a fixed-effects meta-analysis of both. An additional local ancestry specific method was used for PRS weighting. Performance, measured as the Pearson’s correlation between the true and GWAS estimated risk score, is shown across 50 simulations. Simulations assume 1000 causal variants and a heritability of 0.5 to compute the true genetic risk. A p-value of 0.01 and LD r^2^ cutoff of 0.2 was used to select variants for the estimated risk scores.

The most accurate PRS approach varied by the proportion of European ancestry. Populations with greater than 20% African ancestry benefited significantly from the inclusion of population specific weights (Figure 2). Intermediate European ancestry benefitted most from using fixed-effects meta-analysis weighting instead of European weights (r = 0.64 vs. 0.61, *p* = 0.02). In contrast, variant weighting from an African ancestry GWAS instead of from European had higher accuracy in low European ancestry (r = 0.65 vs. 0.53, *p* = 0.009) and African-only (r = 0.64 vs. 0.45, *p* = 2.02×10^−44^) populations. Individuals with high European ancestry had similar accuracy with weights from a fixed-effects meta-analysis as from European (r = 0.73 in both, *p* = 0.79), but decreased performance with the inclusion of weights from an African ancestry GWAS (r = 0.62 vs. 0.73, *p* = 0.01).

No clear benefits, and in some cases significant decreases, were observed for local ancestry informed weights compared to weights from a European or African ancestry GWAS or fixed-effects meta-analysis. Individuals with high, intermediate, and low European ancestry had lower accuracy using local ancestry informed weights compared to the best weighting in each ancestry group: r = 0.71 vs. 0.73 (from fixed-effect or European weights; *p* = 0.58); r = 0.61 vs. 0.64 (from fixed-effect weights; *p* = 0.004); and r = 0.63 vs. 0.65 (from African weights; *p* = 0.60), respectively (Figure 2).

### Performance of Non-European PRS Variant Selection and Weighting Approaches

In our simulations, population specific weighting of PRS SNPs discovered in European ancestry populations improved PRS accuracy; however, the disparity between performance in European ancestry individuals versus African and admixed ancestry individuals remained large. We aimed to explore the potential improvements in PRS that could be gained by including variants discovered in large, adequately powered African ancestry cohorts. Following clumping and thresholding of significant variants using LD and summary statistics from the simulated African populations, an average of 5269 (range = [4462-6071]) variants were included in the PRS (Table 1) reflective of the greater genetic diversity and decreased LD compared to Europeans^23^. In contrast, when constructing a PRS using the same LD and p-value criteria applied to a fixed-effects meta-analysis of European and African ancestry, an average of only 92 (range = [38-197]) variants were included in the PRS. This substantially smaller number was a result of few variants being statistically significant in both populations. Of the total number of variants included from the European GWAS, African ancestry GWAS, and fixed-effects meta, only 1.15%, 0.54%, and 15.0% on average were the exact causal variant from the simulation; an additional 3.72%, 5.34%, and 33.3% tagged at least one causal variant with r^2^ > 0.2 (and were within ±1000 kb of that causal variant) in European ancestry populations and 3.45%, 2.42%, and 28.1% in African ancestry populations (Table 1). The limited overlap between true causal and GWAS selected variants is a result of causal variants in our simulation arising from the total spectrum of allele frequencies, and therefore more likely to be rare, while GWAS is better powered to detect common variants in the study population from which they were identified^2^. These common variants may not adequately tag rare variants due to low correlation^24^.

Overall, we constructed twelve PRS with variants selected from GWAS in European or African ancestry populations or a fixed-effects meta of both (three scenarios) and weights from the same approaches plus an additional local ancestry specific weighting method (four scenarios) (Figure 2). For Europeans, the highest PRS accuracy was achieved with European selected variants and weights (r = 0.77; 95% CI = [0.76, 0.77]); however, a similar accuracy was observed for weights from a fixed-effects meta (r = 0.76; p = 0.53). For Africans, the highest PRS accuracy was with African selected variants and weights from a fixed-effects meta (r = 0.75; 95% CI = [0.74, 0.75]), similar performance was observed with African variants and weights (r = 0.74, p = 0.28). For admixed individuals, the highest performing PRS depended on the population ancestry percentage. In individuals with high European ancestry (>80%), the best PRS was with European selected variants and fixed-effects meta or European weights (r = 0.73; 95% CI = [0.72, 0.74]). For individuals with intermediate (20%-80%) or low (<20%) European ancestry, the most accurate PRS was from using African selected variants and weights from a fixed-effects meta-analysis (r = 0.68; 95% CI = [0.67, 0.68] and 0.71; 95% CI = [0.70, 0.72], respectively). Again, no benefit was observed with the inclusion of local ancestry specific weights for any set of PRS variants. Using a more stringent p-value threshold and including fewer variants into the PRS did not result in a change of the best PRS variants and weights (Figure S2).

### Inclusion of Variants from Diverse Populations

We found that including in the PRS variants discovered in African ancestry GWAS with population specific weights results in less disparity in PRS accuracy across ancestries compared to European selected variants, confirming that GWAS in non-bottlenecked populations may yield a more unbiased set of disease variants^25^. For example, applying to individuals of African ancestry a PRS derived from GWAS variants and weights discovered in training data from the target population results in a 15.7% higher accuracy compared to using a PRS comprised of variants discovered in a European GWAS (also with African weights). In contrast, the gains in accuracy achieved by sourcing variants from ancestry-matched studies were much lower in European ancestry individuals. Compared to a PRS with variants from an African ancestry GWAS (with European weights), a PRS derived from a European GWAS (also with European weights) only gave a 3.9% higher accuracy. We also observed better generalization of PRS based on African selected variants across all admixed groups (Figure 2).

Unlike in Europeans, a PRS with variants discovered in African ancestry populations generalized across ancestral groups with population-specific weighting. However, similar to the European PRS, the African ancestry derived PRS (with African variants and weights) was estimated to have a 1.62% increase in the variance explained of the total trait liability by the PRS for each 10% increase in African ancestry (Figure S4). Through a linear combination of the European and African ancestry derived PRS (Methods)^10^, the relationship between ancestry and accuracy diminished to less than a 0.4% increase per 10% increase of African ancestry (Figure S4).

While the best single PRS for admixed individuals with at least 20% African ancestry selected variants based on a GWAS in an African ancestry population with weights from a fixed-effects meta-analysis, a linear combination of the European and African ancestry derived PRS had higher accuracy; this was particularly true at decreased African ancestry cohort sizes. We saw considerable improvements with the combined PRS over using a European derived (European selected variants and weights) PRS, especially for low European ancestry (CEU < 20%) where even with 10-fold fewer African samples there was a 27.4% increase in PRS accuracy compared to the European derived risk score and a 12.3% increase compared to a PRS with African ancestry selected variants and weights from a fixed-effects meta (Figure 3).

**Figure 3.**
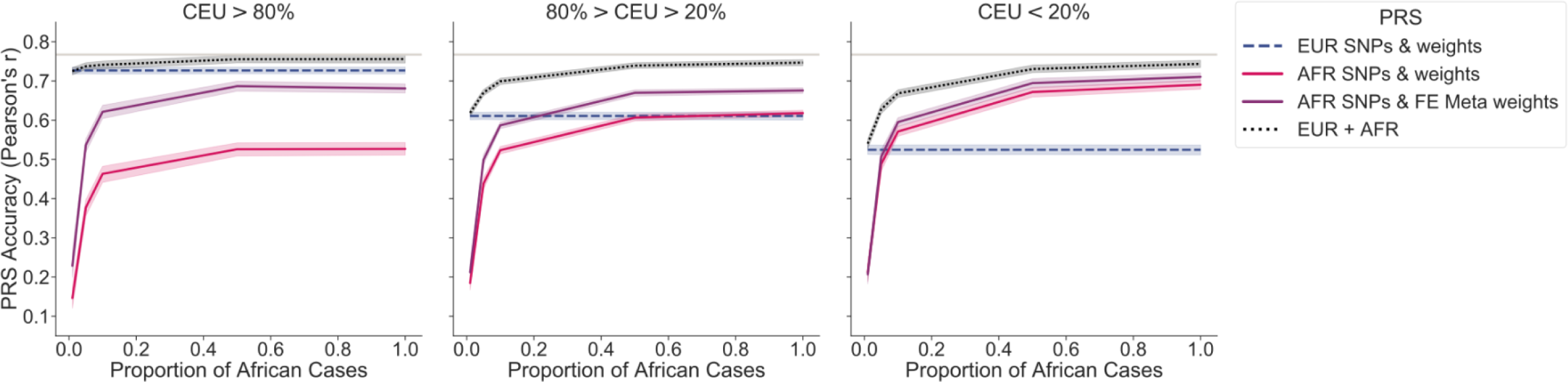
Impact of African Sample Size on PRS Accuracy and Generalization. **PRS accuracy in diverse populations can be improved by including data from an African Ancestry GWAS with smaller sample sizes than in a European GWAS**. The number of African samples used in the GWAS and subsequent PRS construction was decreased to reflect availability of diverse samples in real data. Analysis was conducted assuming 1%, 5%, 10%, 50%, and 100% (matched size of European dataset) of the total African ancestry cases. Average accuracy and the 95% confidence interval were reported across the 50 simulations for different variant selection and weighting approaches. Simulations assume 1000 causal variants and a heritability of 0.5 to compute the true genetic risk. A p-value of 0.01 and LD r^2^ cutoff of 0.2 was used to select variants for the estimated risk score. A linear mixture of single population PRS (*α*_1_*EUR* + *α*_2_*AFR*), with variants and weights selected from that population, was also tested in the admixed population. The mixture coefficients (*α*_1_ and *α*_2_) were estimated in an independent African ancestry testing population.

### Allele Frequency and Linkage Disequilibrium of GWAS variants

We sought to understand what factors impacted PRS generalizability across the different variant selection approaches. GWAS performed in European and African ancestry populations (for SNPs with MAF ≥ 0.01) were both more likely to identify significant variants that were more common in their own population than in the other. Approximately 60% of variants identified in European ancestry populations had minor allele frequencies less than 1% in African ancestry populations and vice-versa; however, given the underlying assumption of homogeneity, the smaller number of variants selected by a meta-analysis of the two populations tended to have more similar minor allele frequencies (Figure 4a). Although European and African ancestry GWAS were both better powered to detect variants at intermediate frequencies within the same study population, GWAS in European ancestry populations may be unable to capture derived risk variants that have remained in Africa, which could be the result of genetic drift, whereas GWAS in African ancestry populations are not subject to this bias^25^.

**Figure 4.**
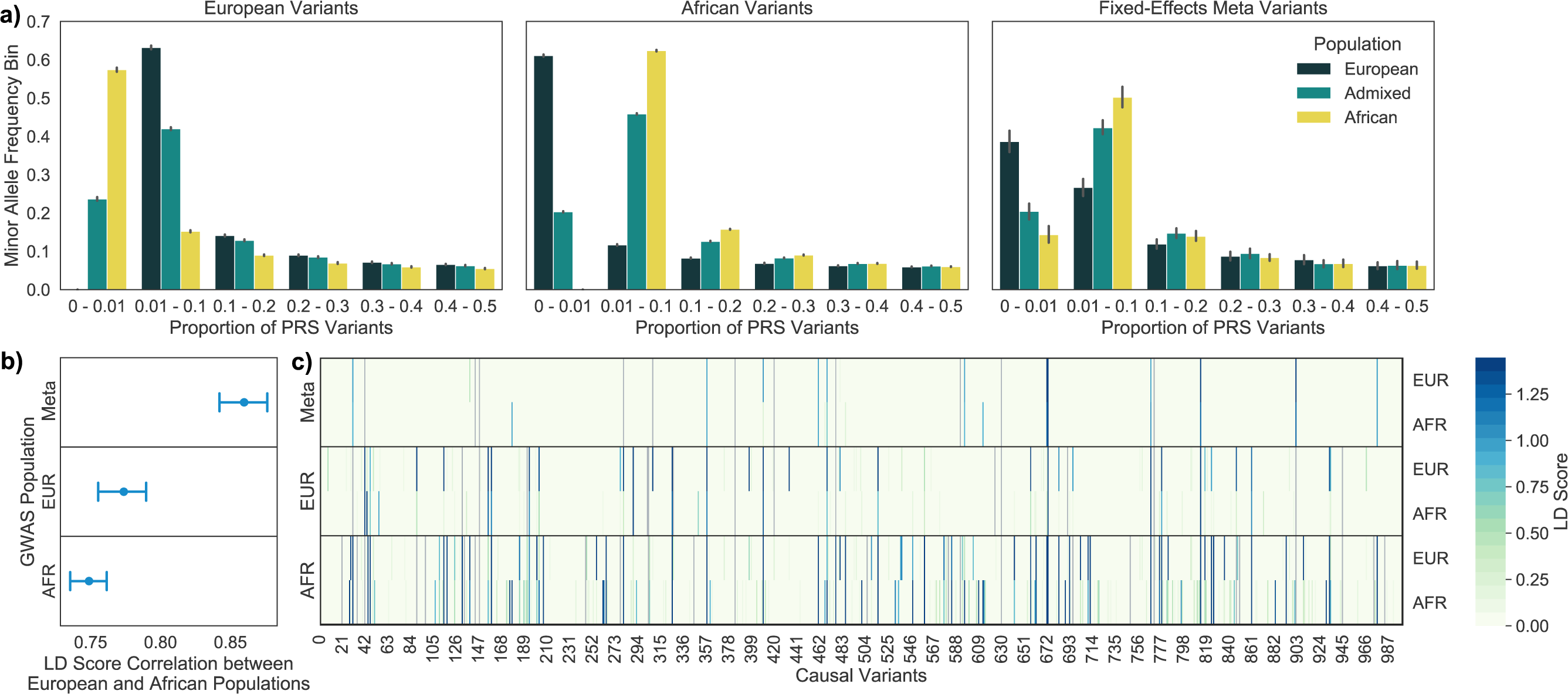
Allele Frequency Distribution of GWAS Selected Variants and LD Tagging of Causal Variants. **GWAS significant variants are more common in the study population from which they were discovered; however, African Ancestry GWAS variants may result in better LD tagging across populations**. Variants were selected from a European or African ancestry GWAS or a fixed-effects meta of both populations. 4a. GWAS variants were binned by their minor allele frequency estimated from the European, African, and admixed populations. The error bar represents the 95% CI across simulations. 4b. LD scores were calculated for every causal variant by adding up the LD r^2^ for each GWAS tag variant within ±1000 kb of the causal variant. LD scores calculated in a Europeans and Africans were compared by Pearson’s correlation. The results were summarized across simulations as the average and 95% CI. 4c. Raw LD scores for each causal variant (m = 1000) calculated in a European or African population for one simulation. Each panel shows the approach used for variant selection. Causal variants directly discovered through the GWAS are colored in grey.

We also examined LD tagging of causal variants by GWAS selected variants within our simulated European and African populations. Each causal variant’s LD score was calculated by summing up the LD r^2^ between that causal variant and every GWAS tag variant within ±1000 kb. The LD scores calculated in European and African ancestry populations were highly correlated (Pearson’s r > 0.7) for the GWAS and fixed-effects meta selected variants. Variants selected from a fixed-effects meta had the highest LD score correlation between populations, as expected given that the variants reached significance in both populations and therefore were more common with similar LD patterns (Figure 4b). Since LD score correlation did not vary largely between simulations, we examined the raw LD scores for a single simulation in order to illustrate differences in LD score magnitude not captured by the Pearson’s correlation.

We found that European selected variants had higher LD scores in European compared to in African ancestry populations; however, variants selected from an African ancestry GWAS tagged causal variants in both populations more strongly (Figure 4c). This is unlikely to be due to the larger number of African selected variants, as the results were unchanged following normalization of LD scores by dividing the total LD score for each causal variant by PRS size (Figure S5). Fixed-effects meta-analysis variants had similar LD score magnitudes. However, while a greater proportion of the fixed-effects meta selected variants were causal, fewer were strong tags for causal variants not directly identified, highlighting the potential need for a model that does not assume homogeneity of effects for tag variants^26^. Additionally, causal variants with identical effect sizes may have differing allele frequencies across populations which would result in heterogeneous allele substitution effects; this helps indicate why a fixed-effects meta-analysis may not be the optimal approach.

### Application to Real Data

To corroborate our simulation findings, we undertook an analysis of 96 PRS developed for the prediction of multiple complex traits in European and African ancestry individuals from the UK Biobank, including HbA1c levels, asthma status, and prostate cancer (Table S1). We tested variant selection strategies based on p-value thresholding and LD clumping of genome-wide summary statistics^15^ computed in independent European or African ancestry cohorts and variant weights from the same approaches with an additional weighting from a fixed-effects meta across populations. Multiple p-value thresholds and weighting strategies were tested to assess the PRS accuracy in African ancestry individuals relative to European ancestry individuals across parameters.

In UK Biobank Europeans, a GWAS significant European-derived PRS (with European variants and weights) had a partial-r^2^ of 1.6%, 1.2%, and 1.5% respectively for HbA1c levels, asthma, and prostate cancer; the same PRS applied to African ancestry individuals, with approximately 13.1% European ancestry^8^, only explained 0.07%, 0.38%, and 0.19% (Figure S6). Although the proportion of variation explained by the PRS (partial-r^2^) was consistently lower in UK Biobank African ancestry individuals compared to Europeans, prediction was improved through the inclusion of variants or weights discovered in an independent African ancestry cohort across all traits (Figure S6). Interestingly, we found that a linear combination of the best performing PRS with European discovered variants and African ancestry discovered variants improved accuracy substantially (Table S2), supporting our simulation finding that a combined PRS which includes variants from the target population, even at smaller sample sizes, is the optimal approach for constructing PRS in admixed and non-European individuals.

## DISCUSSION

Our work shows that incorporating variants selected from European GWAS into a PRS can result in less accurate prediction in non-European and admixed populations in comparison to variants selected from a well-powered African ancestry GWAS. Through simulations and application to real data analysis of multiple complex traits, we provide empirical evidence that supports the use of a linear mixture of multiple population derived PRS to remove bias with ancestry and achieve higher accuracy in admixed individuals with currently available non-European sample sizes. We also demonstrate the anticipated improvements in PRS prediction accuracy that may be achieved with the inclusion of diverse individuals in GWAS, highlighting the need to actively recruit non-European populations.

Our simulation finding that prediction accuracy of a European derived PRS linearly decreases with increasing proportion of African ancestry in admixed African and European populations is consistent with a recent study of height where there was a 1.3% decrease for each 10% increase in African ancestry^8^. This decrease in prediction accuracy has been attributed to linkage disequilibrium and allele frequency differences, as well as differences in effect sizes across populations contributing to height^8^. Our work adds further insights into this reduction in PRS accuracy, showing that (1) it exists in the absence of trans-ancestry effect size differences consistent with previous theoretical models that did look at admixture^2,5^, and (2) variants selected from an African population may not have these same biases. Recent work found that known GWAS loci discovered in Europeans have allele frequencies that are upwardly biased by 1.15% in African ancestry populations which results in a misestimated PRS; a phenomenon that likely arises due to population bottlenecks and ascertainment bias from GWAS arrays^25^. In our simulation study, which was not impacted by ascertainment bias, we show that GWAS in African ancestry populations also identify variants with population allele frequency differences; however, these variants have more consistent LD tagging of causal variants across populations. Our observations support the hypothesis that well-powered African ancestry GWAS will be able to more accurately capture disease associated loci that are more broadly applicable across populations, due to having undergone less genetic drift^25^.

A major advantage of our study is having large simulated European and African ancestry cohorts to provide guidelines for developing the best possible PRS in admixed individuals with current and future GWAS. Through our exploration of 12 PRS, with various variant selection and weighting approaches, we re-capitulate recent results applying similar PRS strategies to an admixed Hispanic/Latino population^9^. For individuals with intermediate proportions of European ancestry (20-80%), we also see improvements using European selected variants and population-specific or fixed-effects meta weights; however, as non-European cohorts get increasingly large it will be imperative to perform variant discovery in these populations as gains in accuracy with weight adjustment of European selected variants will be limited especially in individuals with higher proportions of non-European ancestry.

Current methods for improving PRS accuracy in diverse populations have prioritized the inclusion of variants from European GWAS, as these have higher statistical power to identify trait associated loci. For example, one such approach uses a two-component linear mixed model to allow for the incorporation of ethnic-specific weights^6^. Another method creates ancestry-specific partial PRS for each individual based on the local ancestry of variants selected from a European GWAS^7^. This approach was found to improve trait predictability, compared to a traditional PRS with population specific or European weights, in East Asians for BMI but not height^7^. In contrast, implementing this local-ancestry method^7^ in our simulation, we found that PRS accuracy was higher with African or fixed-effects meta weighting than with local ancestry in admixed African ancestry populations. Our results suggest that true equality in performance will be difficult to obtain solely based on European-identified variants even with local ancestry-adjusted weights. Although local ancestry weighting may have greater benefits when complete sharing across populations is not assumed, we show that in the absence of population-specific factors, the optimal PRS approach involves using variants identified in a large African population and population-specific weighting.

To create a multi-ancestry PRS without incorporating local ancestry, *Márquez-Luna et al. (2017)* uses a mixture of PRS taking advantage of existing well-powered GWAS studies and supplementing with additional information that can be gained from a smaller study in the population of interest^10^. While this approach may offer relative improvement in PRS accuracy for non-Europeans compared to a European-derived PRS, our simulation suggests that the inclusion of significant tag variants discovered in Europeans may unnecessarily hinder predictive performance in non-Europeans. We investigate this approach in the context of varying admixture proportions and find that it achieved high accuracy across all admixed individuals, was not biased by ancestry, and significantly improved performance over a European-only PRS with 10-fold fewer African ancestry cases. Thus, a combination of multiple single population PRS may be the best currently available approach for admixed individuals, and this approach will likely continue to improve as the individual PRS are further developed.

An important novel finding of our work that the inclusion of variants from an African-ancestry population results in less disparity in PRS accuracy across other populations, illustrates the need to recruit diverse populations in GWAS and make these data readily available. Large consortia such as H3Africa, PAGE, the Million Veterans Program, and All of Us are undertaking efforts to support this initiative. Based on our analysis of HbA1c, asthma, and prostate cancer in the UK Biobank, we find that improvement in PRS prediction accuracy is currently possible by incorporating findings from GWAS in African ancestry populations, albeit with lower power. In addition to smaller sample sizes, this potential improvement may be limited by ascertainment bias in what SNPs are included on genotyping arrays and poorer imputation in non-Europeans. GWAS arrays and their imputation have substantially higher coverage among Europeans, and this may result in decreased PRS portability of findings across traits; in such situations, whole genome sequencing in diverse populations may be needed in order to improve accuracy^27,28^. Our study and others support the immense genetic diversity that can be unlocked by studying underrepresented populations to both eliminate the disparity in genetics for prediction medicine and provide novel insights into disease biology for all populations^25,27,29^.

Although our simulation study provides important insight into the future of PRS use, it has important limitations. First, while coalescent simulations allow for decreased computational burden, model assumptions may result in inaccurate long-range linkage disequilibrium especially for whole genome simulations^30^. However, given we only simulated chromosome 20, biases are expected to be modest^30^. We also use a case-control framework for our simulation; therefore, power and potential differences in population PRS accuracy may be even higher if a quantitative trait was used. In addition, our simulations assume random mating among admixed individuals and therefore do not reflect the more complex assortative mating that may be observed, which may impact the distribution of local ancestry tract lengths in our simulation and therefore hinder the improvement of PRS accuracy by local ancestry weighting^31^. Also, although we provide evidence to suggest the contribution of population differences in allele frequency and LD tagging of causal variants to loss of PRS accuracy with varying ancestry, we do not delineate how each of these factors decrease accuracy independently; this is a direction for future work. Finally, we have only simulated individuals from Yoruba, a West African population, which is not representative of the greater diversity in Sub Saharan Africa^32^. Future studies must be done to ensure our findings can be extended to individuals from other regions of Africa.

Overall, our findings support the concern that while studies have demonstrated the potential clinical utility of PRS, adopting the current versions of these scores could contribute to inequality in healthcare^4^. Going forward, future studies should prioritize the inclusion of diverse participants and care must be taken with the interpretation of currently available risk scores. While statistical approaches may offer improvements in accuracy compared to current European-derived risk scores, in order to truly diminish the disparity and achieve PRS accuracies similar to in European ancestry populations we must actively recruit and study diverse populations.

## SUPPLEMENTAL DATA

Document S1. Figures S1-S6 and Tables S1-S2

## ACKNOWLEDGEMENTS

This material is based upon work supported by the National Science Foundation Graduate Research Fellowship Program under Grant No. 1650113 and NIH grant CA201358. Any opinions, findings, and conclusions or recommendations expressed in this material are those of the author(s) and do not necessarily reflect the views of the National Science Foundation. This research has been conducted using the UK Biobank Resource under Application Number 14015. Furthermore, the authors thank Linda Kachuri for providing helpful feedback and discussion.

## WEB RESOUCES

HBA1 summary statistics (*Wheeler et al*. 2018): https://www.magicinvestigators.org/downloads/

Asthma summary statistics (*Daya et al. 2019* and *Demenais et al. 2018*): https://www.ebi.ac.uk/gwas/downloads/summary-statistics

PrCa summary statistics (Emami et al. 2020): https://www.ncbi.nlm.nih.gov/projects/gap/cgi-bin/study.cgi?study_id=phs001221.v1.p1

plink2: https://www.cog-genomics.org/plink/2.0/

RFMix: https://github.com/slowkoni/rfmix

METASOFT: http://genetics.cs.ucla.edu/meta_jemdoc/

## DATA AND CODE AVAILABILITY

The code generated during this study is available at https://github.com/taylorcavazos/PRS_Admixture_Simulation

