## Supplementary Data for "Inclusion of Variants Discovered from Diverse Populations Improves Polygenic Risk Score Transferability"

### SUPPLEMENTAL MATERIALS

**Figure S1.** European Derived Risk Score Accuracy with Varying Simulation Causal Variants and Assumed Heritability

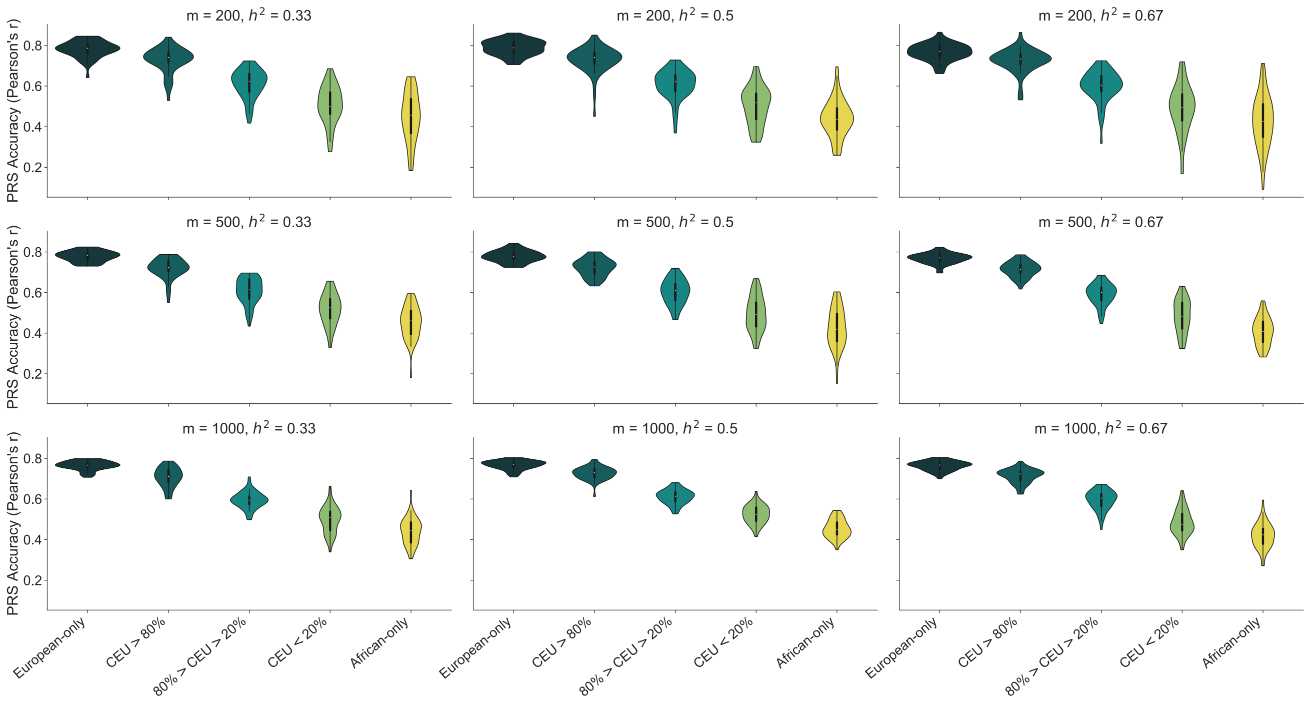


**Figure S1 Legend:** Our simulation assumes a set number of causal variants (m) and trait heritability ($h^2$) when generating the true genetic risk score. We varied these parameters and tested all combinations assuming m = {200, 500, 1000} and $h^2$ = {0.33, 0.5, 0.67}. The accuracy was measured by Pearson's correlation between the true and GWAS estimated risk score for each of the 50 simulations. We used a European GWAS to select independent variants and effect sizes for the PRS ($p < 0.01$ and LD $r^2 = 0.2$). This risk score was applied to Europeans, Africans, and admixed populations.

**Figure S2.** PRS Accuracy with Varying P-value Thresholds

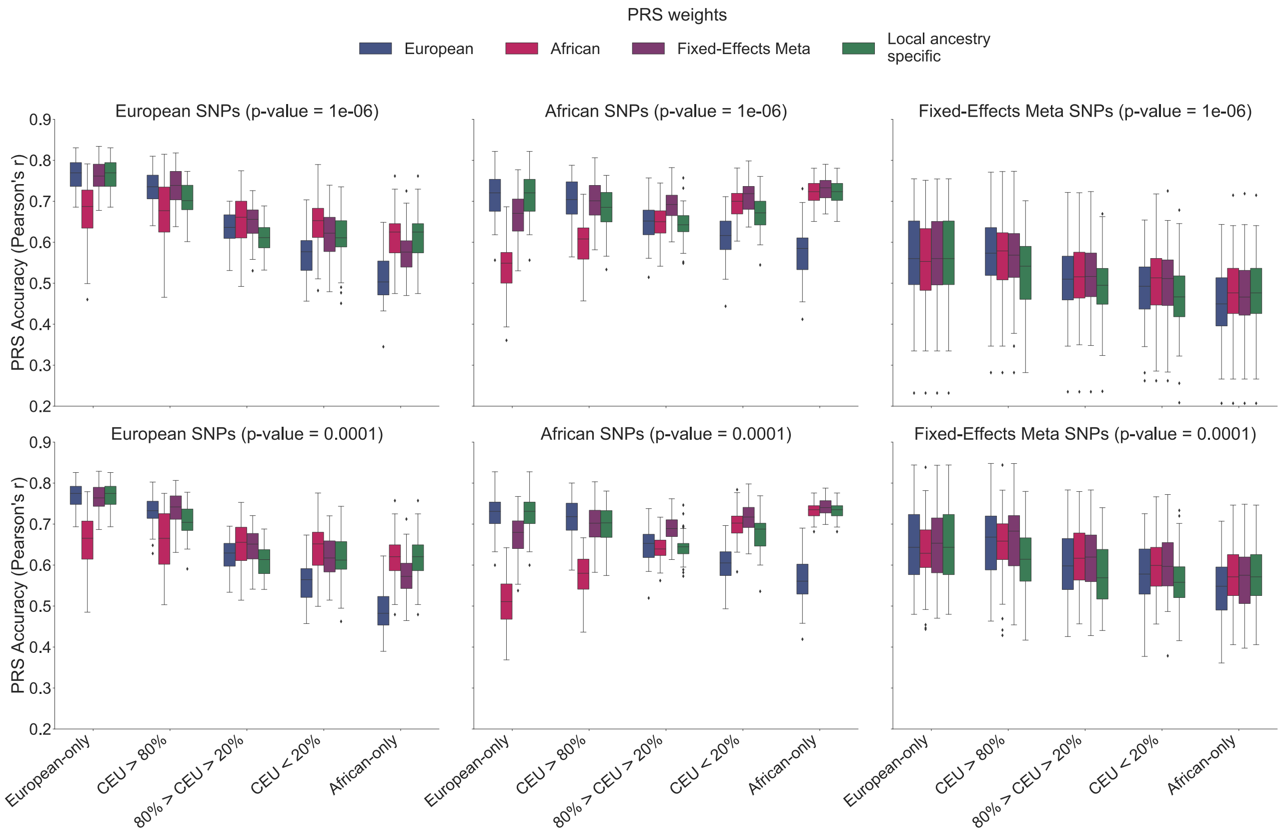


**Figure S2 Legend:** The p-value used for variant selection approach was decreased to allow fewer variants into the PRS. In addition to $p < 0.01$, shown in the main text results, we also tested $p < 1x10^{-4}$ and $1x10^{-6}$. Simulations assume 1000 causal variants and a heritability of 0.5 to compute the true genetic risk. A p-value of 0.01 and LD $r^2$ cutoff of 0.2 was used to select variants for the estimated risk scores.

**Figure S3.** Total Liability Variance Explained by a European Derived Risk Score as a Function of Genetic Admixture with Varying Simulation Parameters

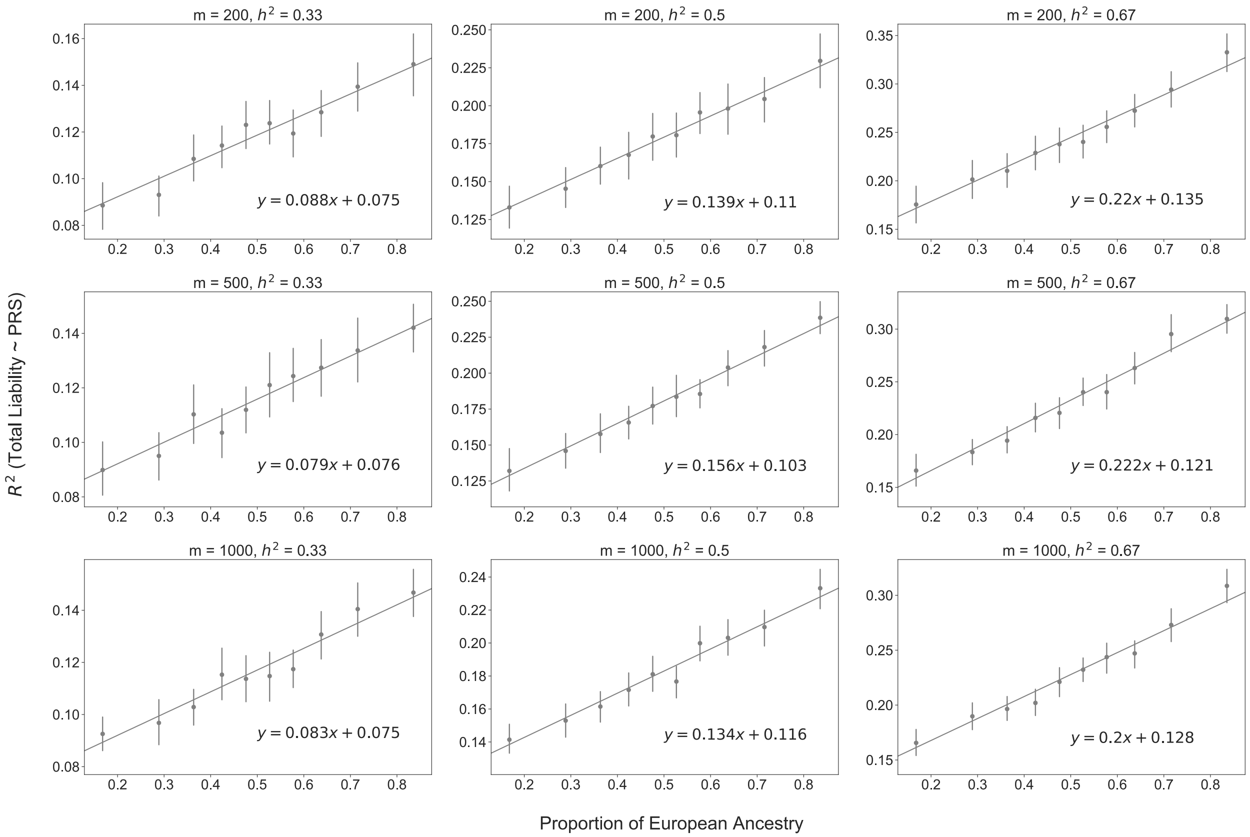


**Figure S3 Legend:** Admixed individuals were split into deciles of global genome-wide European ancestry. For each simulation trial the variance explained ($R^2$) of the total trait liability (genetics + environment) by the estimated PRS (European variants and weights) was computed within each ancestry decile. Across simulations the average ancestry for each bin is reflected as the point and the vertical bar represents the 95% confidence intervals. The equation shows the slope of each line.

**Figure S4:** Linear Mixture of Multiple Population Derived PRS Removes Bias Between Accuracy and Ancestry

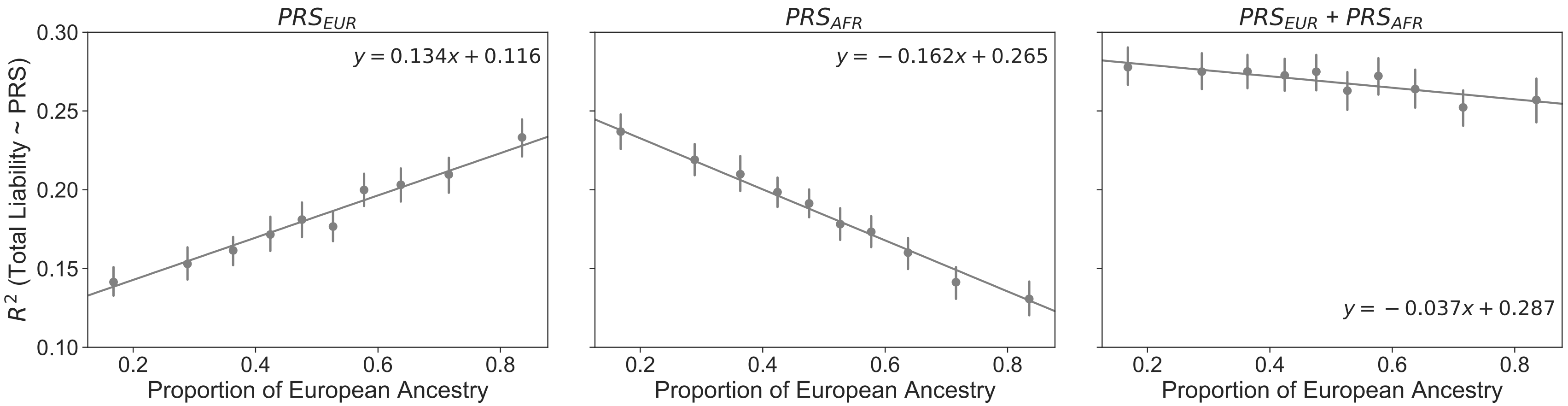


**Figure S4 Legend:** Two population derived risk scores ($PRS_{EUR}$ and $PRS_{AFR}$) were constructed from variants and weights from GWAS in the designated simulated population and applied to an independent simulated admixed population. The PRS were combined ($\alpha_1 PRS_{EUR} + \alpha_2 PRS_{AFR}$) through a linear mixture approach described by Márquez-Luna et. al.[1] where the mixing coefficients ($\alpha_1$ and $\alpha_2$) were estimated in an independent African ancestry testing population and validated in the admixed population. Admixed individuals were split into deciles of global genome-wide European ancestry and for each simulation trial variance explained ($R^2$) of the total trait liability by each PRS approach was calculated within each decile. Across simulations the average ancestry for each bin is reflected as the point and the vertical bar represents the 95% confidence intervals. The equation shows the slope of each line.

**Figure S5.** Normalized LD Tagging of Causal Variants by GWAS Selected Variants

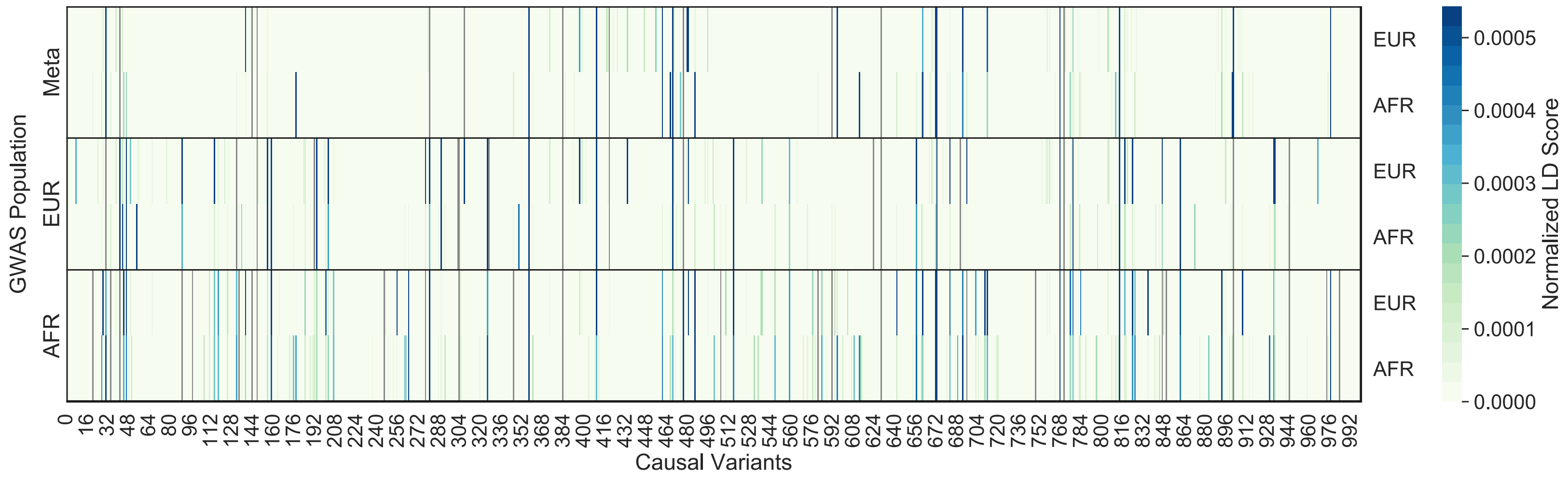


**Figure S5 Legend:** LD scores were calculated for every causal variant by adding up the LD $r^2$ for each GWAS tag variant within ±1000 kb of the causal variant. LD scores were then normalized by the total number of GWAS selected variants from each population reflected by the 3 panels. The normalized LD scores for each causal variant (m = 1000) calculated in a European or African population is shown for one simulation. Causal variants directly discovered through the GWAS are colored in gr

**Figure S6:** PRS Performance Across SNP Selection and Weighting Approaches for Multiple Complex Traits in the UK Biobank

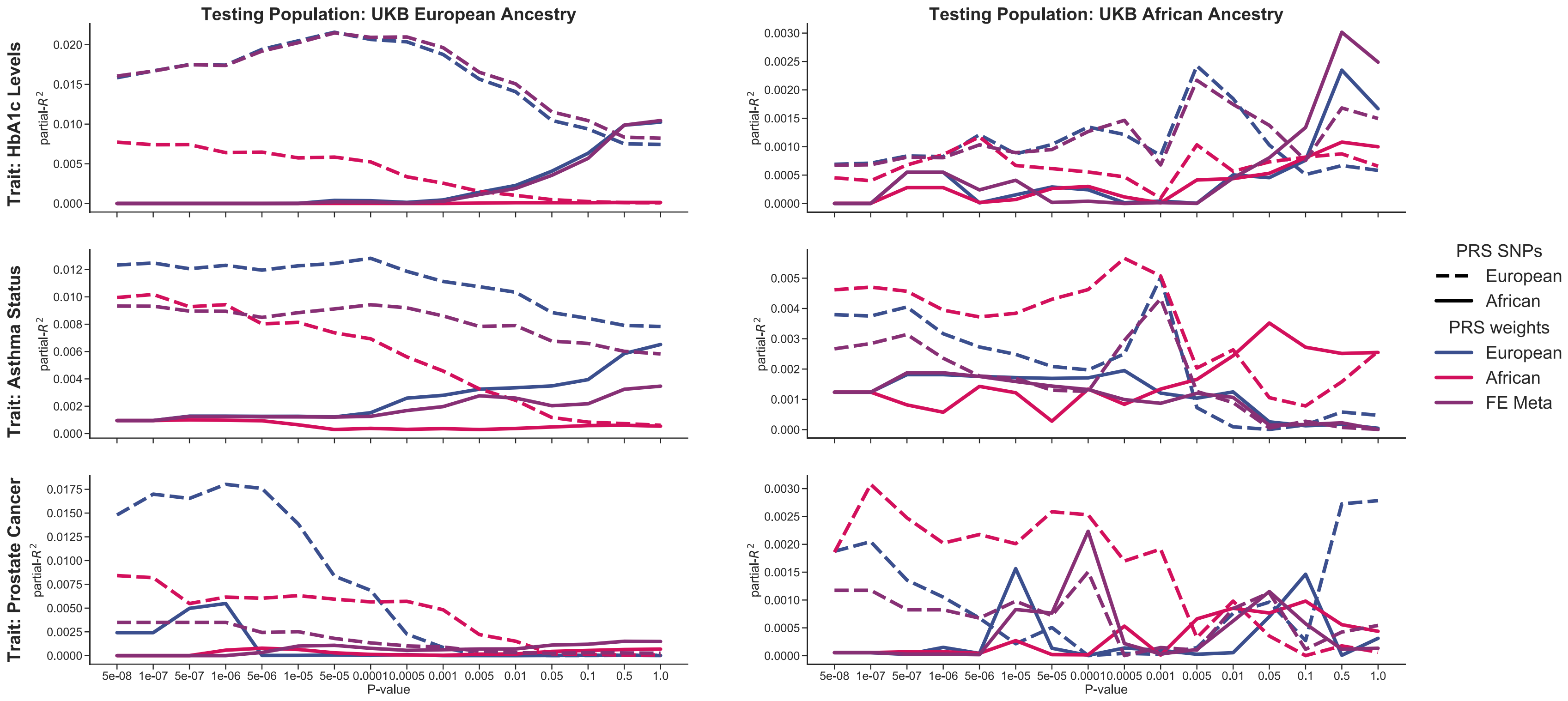


**Figure S6 Legend:** Partial-$R^2$ was calculated for each PRS, within Europeans and Africans from the UK Biobank, by subtracting the variance explained by the null model. For HbA1c we fit a linear model with age, sex, BMI, and PCs1-10 with and without the PRS. Similarly for asthma status and prostate cancer, we determined the Nagelkerke's pseudo partial-R2 following logistic regression of case status on age, sex (asthma only), BMI (prostate cancer only), and PCs1-10 with and without the PRS.

**Table S1.** Independent GWAS Summary Statistics and UK Biobank Testing Population used for PRS

| Trait | Study | # Variants | # Samples Summary Statistics (cases) | | # Samples Testing – UK Biobank (cases) | |
|---|---|---|---|---|---|---|
| | | | European Ancestry | African Ancestry | European Ancestry | African Ancestry |
| HbA1c | Wheeler et al. 2017[2] | 1,768,940 | 123,665 (NA) | - | 395,472 (NA) | 5,886 (NA) |
| | Wheeler et al. 2017[2] | 1,545,588 | - | 7,564 (NA) | | |
| Asthma | Demenais et al. 2018[3] | 1,202,829 | 127,669 (19,954) | – | 413,870 (54,537) | 7,250 (1,002) |
| | Daya et al. 2019[4] | 12,955,021 | – | 14,654 (7,009) | | |
| Prostate Cancer | Emami et al. 2020[5] (KP RPGEH) | 21,230,454 | 11,649 (6,196) | – | 413,870 (7,610) | 7,250 (166) |
| | Conti et al. 2020[6] (AAPC) | 9,910,837 | – | 9,531(4,853) | | |

**Table S1 Legend:** Description of cohort sizes of independent summary statistics used for PRS construction and UK Biobank population used for assessing PRS accuracy. Case counts are not applicable (NA) for non-binary traits. For prostate cancer, GWAS studies included multiple populations; however, we only used the Kaiser Permanente Research Program in Genes, Environment, and Health (KP RPGEH) summary statistics from *Emami et al. 2020* and the African American Prostate Cancer (AAPC) consortium data from *Conti et al. 2020*. Number of variants from each independent study that were available for analysis in both testing populations from the UK Biobank are reported here.

**Table S2.** Cross-Validated Accuracy for a Linear Combination of Multiple PRS in the UKB African Ancestry Population

| Trait | Best PRS – African Selected SNPs | | | | Best PRS – European Selected SNPs | | | | Combined PRS |
|---|---|---|---|---|---|---|---|---|---|
| | Weights | P-value | # SNPs | Partial-$R^2$ | Weights | P-value | # SNPs | Partial-$R^2$ | Partial-$R^2$ |
| HbA1c | FE Meta | 0.5 | 146,670 | 0.0040 | European | $5x10^{-3}$ | 2,538 | 0.0022 | 0.0052 |
| Asthma | African | 0.05 | 157,573 | 0.0036 | African | $5x10^{-4}$ | 556 | 0.0060 | 0.0089 |
| Prostate Cancer | FE Meta | $1x10^{-4}$ | 190 | 0.0047 | African | $1x10^{-7}$ | 10 | 0.0082 | 0.0129 |

**Table S2 Legend:** The PRS with the highest Partial-$R^2$ in the full UKB African ancestry testing cohort was chosen using African ancestry or European ancestry selected SNPs from independent summary statistics for each trait. A combined PRS was constructed by using a linear combination of the two best PRS ($\alpha_1 PRS_{EUR} + \alpha_2 PRS_{AFR}$). Through 5-fold cross validation, 80% of the cohort was used to estimate the mixing coefficients ($\alpha_1$ and $\alpha_2$) and the accuracy of combined PRS was tested in the remaining 20% of the cohort. For disease traits, stratified 5-fold cross validation was used to ensure a consistent ratio of cases and controls across folds. Accuracy was reported as the partial-$R^2$ or the proportion of variation explained by the PRS. Accuracy of the single PRS were also re-assessed through 5-fold cross validation for comparison.
